# Morphological profiling of tubercule bacilli identifies drug pathways of action

**DOI:** 10.1101/2020.03.11.987545

**Authors:** Trever C. Smith, Krista M. Pullen, Michaela C. Olson, Morgan E. McNellis, Ian Richardson, Sophia Hu, Jonah Larkins-Ford, Xin Wang, Joel S. Freundlich, D. Michael Ando, Bree B. Aldridge

## Abstract

Morphological profiling is a method to classify target pathways of antibacterials based on how bacteria respond to treatment through changes to cellular shape and spatial organization. Here, we utilized the cell-to-cell variation in morphological features of *Mycobacterium tuberculosis* bacilli to develop a rapid profiling platform called <u>Morph</u>ological <u>E</u>valuation and <u>U</u>nderstanding of <u>S</u>tress (MorphEUS). MorphEUS classified 94% of tested drugs correctly into broad categories according to modes of action previously identified in the literature. In the other 6%, MorphEUS pointed to key off-target or secondary bactericidal activities. We observed cell-wall damaging activity induced by bedaquiline and moxifloxacin through secondary effects downstream from their main target pathways. We implemented MorphEUS to correctly classify three compounds in a blinded study and identified an off-target effect for one compound that was not readily apparent in previous studies. We anticipate that the ability of MorphEUS to rapidly identify pathways of drug action and the proximal cause of bactericidal activity in tubercule bacilli will make it applicable to other pathogens and cell types where morphological responses are subtle and heterogeneous.

**Significance Statement:** Tuberculosis is a leading cause of death in the world and requires treatment with an arduous multidrug regimen. Many new tuberculosis drugs are in development, and the drug development pipeline would benefit from more rapid methods to learn drug mechanism of action and off-target effects. Here, we describe a high throughput imaging method for rapidly classifying drugs into categories based on the primary and secondary cellular damage called <u>Morph</u>ological <u>E</u>valuation and <u>U</u>nderstanding of drug-<u>S</u>tress (MorphEUS). We anticipate that MorphEUS will assist in rapidly pinpointing causes of cellular death in response to drug treatment in tuberculosis and other bacterial pathogens.

## Introduction

*Mycobacterium tuberculosis* (Mtb), the causative agent of tuberculosis (TB), causes more deaths annually than any other infectious agent (1). Tuberculosis treatment is lengthy, lasting between four months to over a year (1). The difficult regimen, rate of relapse, and incidence of drug resistant Mtb has motivated a significant effort to develop new antibacterial compounds that are effective in sterilizing Mtb infection (2). Many new drug classes and derivative compounds have been developed (2), but rapidly identifying the primary and secondary pathways of action of each compound is often a protracted process due to the difficulty in generating resistant mutants, and dissecting the broad-reaching metabolic effects of drug treatment leading to death (3). Furthermore, drug action on bacterial cells can elicit dynamic responses in multiple pathways both on- and off-target, some of which are specific to bacterial growth environment and treatment dose (4–6). An approach that combines computational algorithms with traditional methods for identifying pathway of action would enable faster drug development.

In other bacterial systems such as *Escherichia coli*, *Bacillus subtilis*, and *Acinetobacter baumannii*, profiling of cytological changes in response to treatment has yielded a rapid and resource-sparing procedure to determine drug mechanism (7–9). This method, known as bacterial cytological profiling (BCP), groups drugs with similar mechanisms of action by clustering profiles of drug-treated bacteria using multivariate analysis methods such as principal component analysis (PCA) (7–9). BCP is efficient and rapid because these cytological features can be automatically derived from high-throughput images of stained, fixed samples.

We hypothesized that BCP could be similarly utilized to map pathways of drug action in Mtb. We found that Mtb morphological response to treatment were subtle and incorporation of cellular variation metrics improved the ability to profile drug action in Mtb. Here, we present an imaging and data analysis pipeline for Mtb that classifies morphological profiles called MorphEUS (<u>Morph</u>ological <u>E</u>valuation and <u>U</u>nderstanding of drug-<u>S</u>tress). We demonstrate that MorphEUS clusters antibacterials by their pathways of action. Because MorphEUS is based on the physical unraveling of cells due to stress, it can be used to identify key proximal cellular stressors that arise from off-target and secondary effects. We used MorphEUS to identify secondary effects for two TB drugs (moxifloxacin and bedaquiline) and a non-commercial compound. We propose that MorphEUS will be useful in classifying drug action in other pathogens where morphological responses are subtle and heterogeneous.

## Results

### Antibacterial treatment induces drug-specific morphological response in mycobacteria

To assess whether mycobacteria exhibit distinct morphological responses to drugs targeting different pathways, we used a chromosome-decorating reporter strain (GFP translational fusion to RpoB) of *Mycobacterium smegmatis* to visualize cell and nucleoid shape characteristics (10) before and during drug treatment by time-lapse imaging. We observed rapid nucleoid condensation in rifampicin-treated *M. smegmatis* that was not apparent in ethambutol-treated cells (Fig.1A). In contrast, moxifloxacin treatment induced filamentation with nucleoid decompression. These antibacterials have different mechanisms of action (inhibitors of transcription, cell-wall synthesis, and DNA synthesis, respectively), providing evidence that the morphological changes induced by drug treatment in mycobacteria are dependent on drug target. These data led us to hypothesize that exploitation of morphological features in mycobacteria would allow clustering of antibacterials in a manner similar to previously described BCP studies.

We next asked whether drug treatment of Mtb, like *M. smegmatis*, elicited well-defined cytological fingerprints. Guided by cytological profiling methods in other bacterial species (7–9), we treated Mtb grown in standard rich growth medium with a high drug dose (3x the 90% inhibitory concentration; IC90) for 17 hours (~1 doubling time). We generated a dataset of morphological features from Mtb treated with 34 antibacterials that encompass a wide range of drug classes and classified the target pathway of each drug according to published findings (Table S1). We imaged fixed, membrane-(FM4-64FX) and nucleoid-(SYTO 24) stained Mtb in biological triplicate. Unlike *M. smegmatis*, *E. coli*, and *B. subtilis*, Mtb did not exhibit striking physical differences that readily distinguish drugs targeting dissimilar cellular pathways (Fig. 1A and Table S1). Furthermore, the morphological responses to drug treatment in Mtb were more subtle than *M. smegmatis* (Fig. 1A). Because Mtb are smaller than *M. smegmatis*, we hypothesized that there may still be quantitatively significant differences among morphological features in drug-treated bacilli that are more sensitive to noise and variation. Using image segmentation and analysis, we quantified 25 morphological features per treatment group (Table S2). We observed significant differences among treatment groups in features such as cell shape, nucleoid shape, and staining intensity (Fig. 1B). However, the resulting morphological profiles from drug treatment did not cluster based on broad drug target categories using PCA (Fig. S1A top).

**Figure 1.**
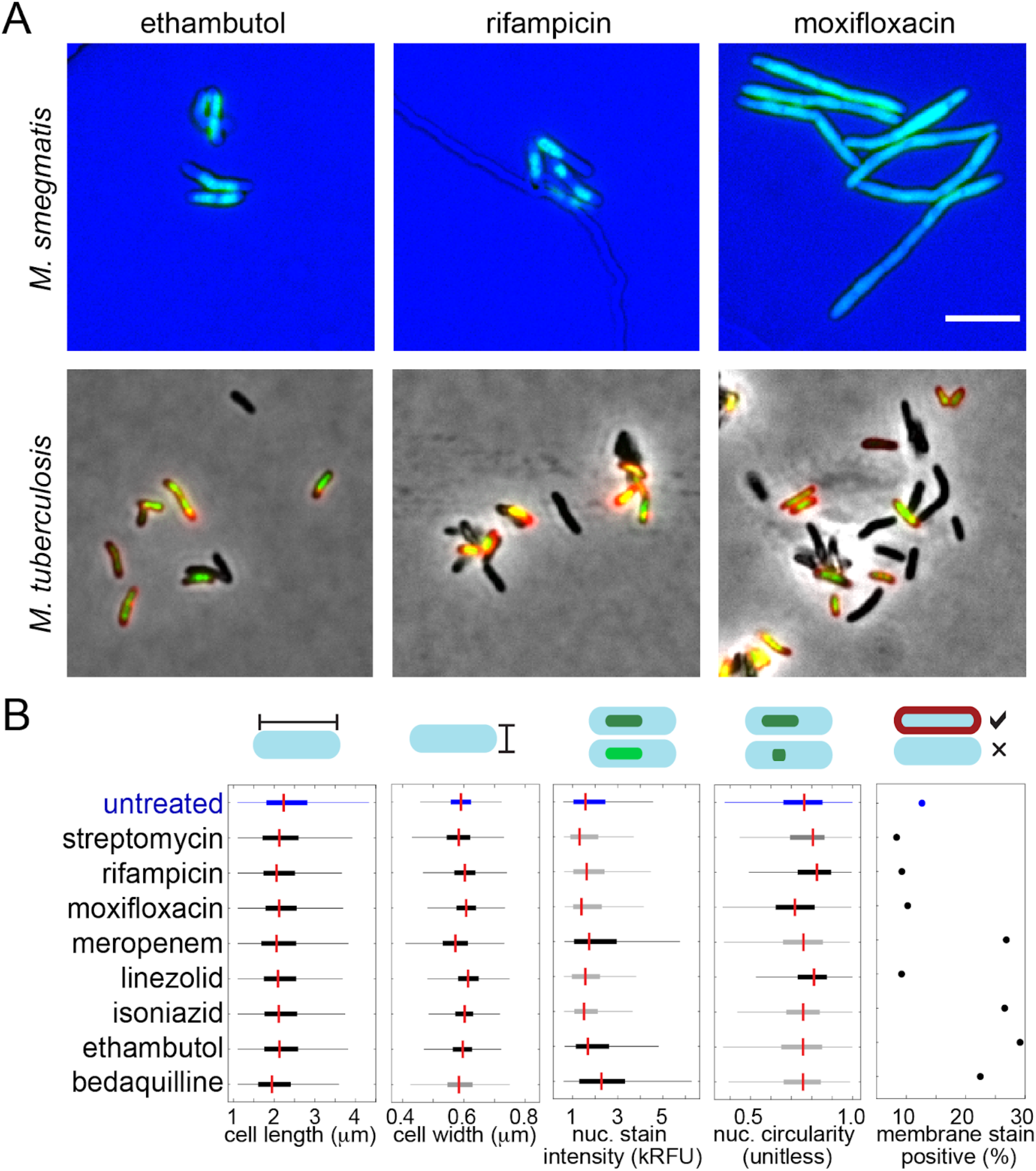
Drug treatment induces distinct morphological characteristics in mycobacteria. (A) Fluorescent time-lapse imaging of *M. smegmatis* (top panels) and fixed-cell imaging of Mtb (bottom panels) treated with ethambutol (left, 3xIC90), rifampicin (middle, 3xIC90) or moxifloxacin (right, 0.25xIC90). Time-lapse images are of an *M. smegmatis* reporter strain expressing RpoB-GFP as a chromosomal marker (green); snapshots are after 6h of treatment. Following 17h treatment, Mtb were PFA-fixed and stained with FM4-64FX (red; membrane) and SYTO 24 (green; DNA). The size bar represents 5 μm. (B) A comparison of select Mtb morphological features across seven antibiotic treatments and untreated control (n=1625-3983). Red lines mark the medians, boxes mark the 25-75th percentiles, and the whiskers extend the range of parameters that are not outliers. Black whiskers and dots indicate p<0.05 compared to untreated control (blue) whereas gray whiskers are not significantly different from untreated using a Kruskal-Wallis test.

One explanation for the poor performance of BCP in Mtb may be the significant cell-to-cell variation in morphological features (Fig. 1 and Fig. S2). This inherent heterogeneity is consistent with the variable nature of Mtb, which on the single-cell level exhibits heterogeneity through asymmetric growth and division, differential drug susceptibility, and metabolic state (11–15). Cell-to-cell variation is most apparent in the ability of Mtb bacilli to take up stains. For example, only ~10% of untreated bacilli are stain positive, whereas the proportion increases to ~30% when treated with cell-wall acting antibacterials (Fig. 1B). We speculated that variation itself was an important feature of drug response that should be captured in the profiling of drug mechanism of action.

### Morphological profiling of Mtb is improved by explicit incorporation of parameters of cellular variation

To capture cell-to-cell variation, we developed a new analysis pipeline for Mtb that incorporates variation as an important class of features to discriminate drug target pathways (Table S1-2, Fig. 2 and Fig. S3, (16)). This analysis formulation also addresses the subtlety of cytological changes by taking into account the full dimensionality of the data to produce discrete classifications. The exploitation of feature variation provided another dimension to distinguish drug categories. For example, when treated with isoniazid, Mtb nucleoid stain intensity was less variable than when treated with bedaquiline or meropenem (Fig. 1B middle). We accounted for a fragile feature selection process (in which several sets of features may achieve similar model accuracy) by performing a series of classification trials (Fig. 2). The resulting analysis was visualized using a network web or matrix describing the frequency of drug-drug links (Fig. 3). Morphological responses were measured in cells treated with high- and low-doses of drugs (Fig. S4) and combined to generate a joint-dose profile to incorporate richer profiling behaviors. We refer to this analysis pipeline as MorphEUS (Morphological Evaluation and Understanding of drug-Stress).

**Figure 2.**
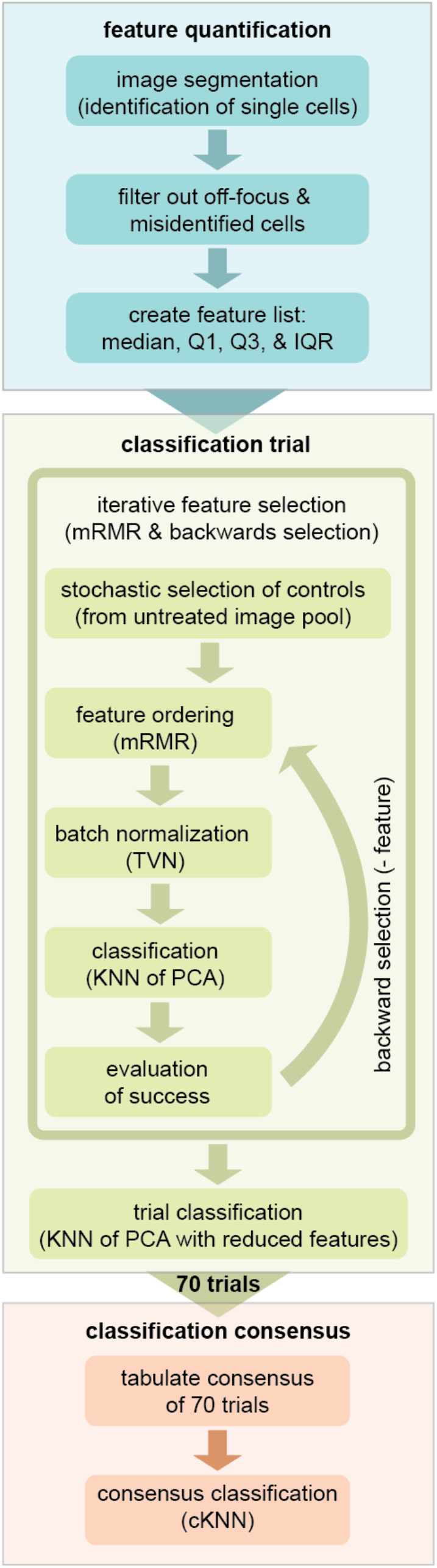
Computational pipeline for MorphEUS. The MorphEUS pipeline is comprised of three into three main steps, feature quantification (blue), classification trials (green), and classification consensus (orange). The main components of each step are highlighted as boxes within each of the three groups. A detailed description of each step is described in the Materials and Methods.

**Figure 3.**
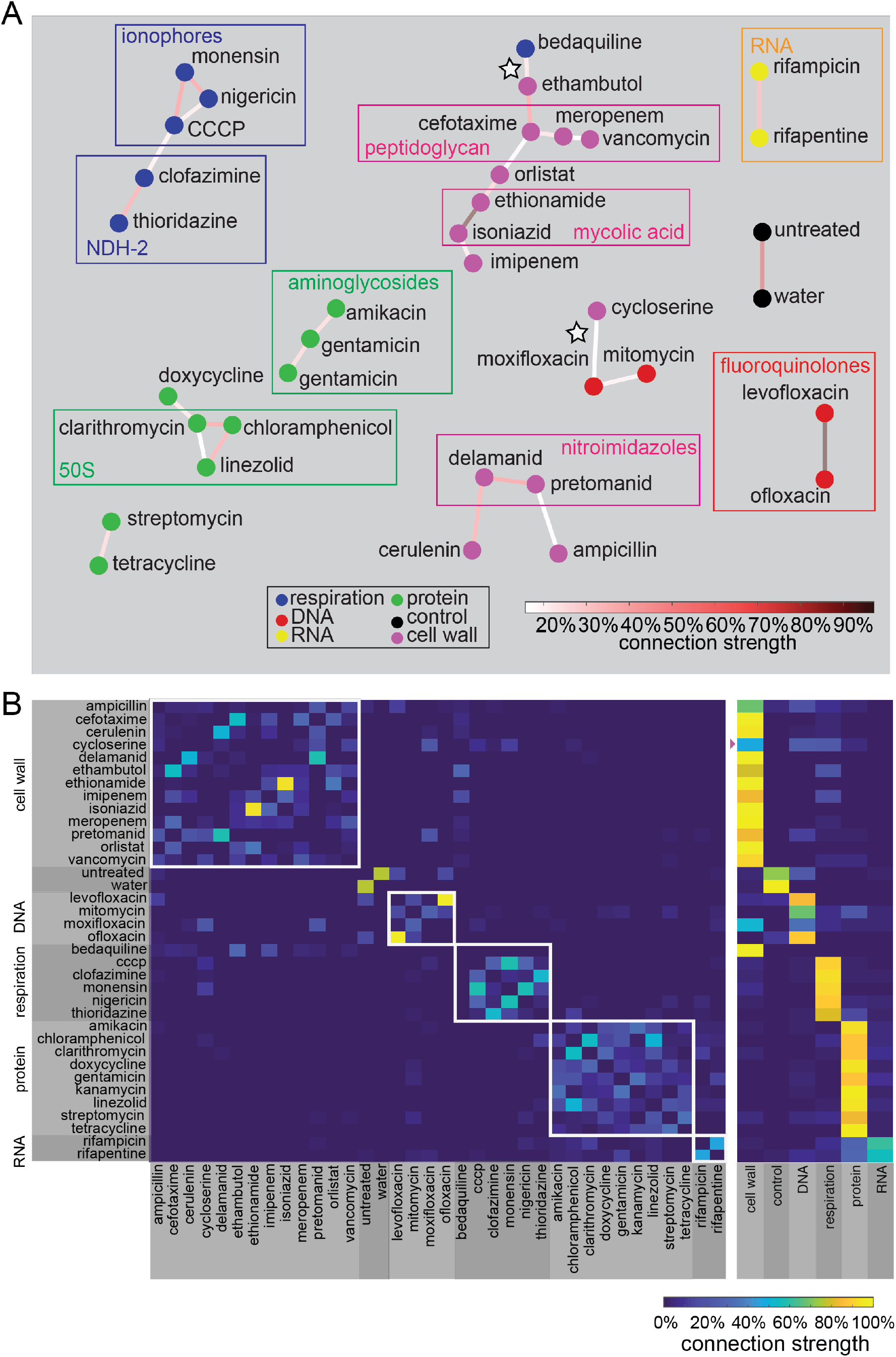
MorphEUS classifies antibacterial compounds by pathway of action. (A) cKNN map of the joint dose profile displaying connections that occur in at least 17% of the classification trials. Drugs within each broad category are represented by nodes of the same color, illustrating whether morphological profiles were similar amongst drugs acting on the same pathway. Rectangles drawn around groups of drugs indicate clustering of drugs that share similar targets within the designated broad category. White stars mark bedaquiline and moxifloxacin, which map to drugs in broad categories other than their own. (B) cKNN matrix of drug nearest neighbor pairings corresponding to (A) by specific drugs (left) and broad categorization (right). The broad drug target categorizations are indicated to the left of the drug names and on the bottom axis of the heatmap on the right. A purple triangle is placed next to the broad categorization for the weakly categorized cell-wall acting drug cycloserine.

Using MorphEUS, drugs in the same broad categories are generally grouped together (94% accurate categorization with 76% accurate cross validation; Fig. 3). Furthermore, within the broad categories, drugs sharing target pathways were found to have stronger connections to one another. For example, within the cell-wall acting category, strong connections were observed between ethionamide and isoniazid (inhibitors of the enzyme InhA), delamanid and pretomanid (nitroimidazole drug class, inhibits the synthesis of mycolic acids), and meropenem, cefotaxime, and vancomycin (all peptidoglycan inhibitors) (Fig. 3 and Table S1). We also observed strong connectivity between inhibitors of cellular respiration with the ionophores CCCP, monensin, and nigericin forming stronger connections with each other compared to clofazimine and thioridazine (both shown to target NDH-2 of the electron transport chain) (17, 18). MorphEUS also predicted strong connections among protein synthesis inhibitors that target the 50S ribosomal subunit (clarithromycin, chloramphenicol, and linezolid). Finally, the fluoroquinolones levofloxacin and ofloxacin grouped together as did rifampicin and rifapentine - inhibitors of transcription.

### Morphological response to treatment reflects key off-target effects

Among the 34 antibacterials profiled, only cycloserine and bedaquiline were miscategorized by general drug group (white stars in Fig. 3A); e.g. their profiles most strongly linked to an antibacterial from a different broad drug category. Cycloserine is a cell-wall acting drug that inhibits the formation of peptidoglycan. Cycloserine weakly profiled with the category of cell-wall acting antibiotics, but its strongest connection to an individual antibacterial was with the fluoroquinolone moxifloxacin (DNA-damaging; white star in Fig. 3). To understand whether MorphEUS correctly predicted the cell-wall targeting activity of cycloserine to be moderate compared to other cell-wall acting drugs, we evaluated the transcriptional profiles of Mtb treated with compounds targeting five different pathways in Mtb using data from previous studies (10, 11) (Fig. S6). We focused on induction of the *iniBAC* operon, which is rapidly upregulated due inhibition of cell-wall synthesis and is used to screen for cell-wall acting compounds (12, 13). We observed that cycloserine’s induction of *iniBAC* genes was mild compared to other cell-wall acting antibiotics (~1.5 fold and ~3.5 fold respectively), consistent with the weak association of cycloserine to other cell-wall antibacterials (Figure 3 Right and Fig S6 Top). We next hypothesized that cycloserine has off-target effects that are DNA damaging and that these effects drive the connection to moxifloxacin. However, we did not find reports of mutants that confer resistance to both moxifloxacin and cycloserine (14–18). An alternative hypothesis is that the connection cycloserine to moxifloxacin is driven by off-target effects by moxifloxacin, such as cell-wall damage. Moxifloxacin’s morphological profiles are highly dose-dependent, resembling the other fluoroquinolones at low-dose but not at high-dose or joint-dose (Fig. S7, Fig 3). This shift away from other fluoroquinolones by moxifloxacin suggests that there is an off-target or secondary effect at high dose.

Transcriptional analysis based on published studies of moxifloxacin-treatment Mtb demonstrated a mild but significant increase in *iniB* expression and a 2.5-fold induction in *ddlA*, a molecular target for cycloserine (Fig. S6 bottom) (16, 17, 19). Taken together these data suggest that the similarity between cycloserine and moxifloxacin morphological profiles arises from an off-target cell-wall damaging effect of moxifloxacin and a mild inhibition of cell-wall synthesis by cycloserine.

The second unexpected profile was for bedaquiline, an ATP-synthesis inhibitor, which mapped to cell-wall acting antibiotics ethambutol and imipenem (Fig. 3A, white star and 3B). Components of the mycobacterial cell-wall, in particular peptidoglycan (PG) and arabinogalactan (AG), are linked to energy production in the cell with components of glycolysis feeding directly into the synthesis of PG and AG (25, 26). In standard laboratory nutrient-replete medium, the presence of sugars allows Mtb to generate ATP from both glycolysis and TCA cycle through substrate-level phosphorylation and oxidative phosphorylation via the electron transport chain (6). Treatment with bedaquiline shuts down the ability of Mtb to carry out oxidative phosphorylation (6) initiating an energy crisis in which Mtb becomes reliant on substrate-level phosphorylation for ATP generation. We hypothesized that bedaquiline disturbed metabolism in a manner that prevents the synthesis of new PG and AG leading to a morphological profile that resembles cells treated with inhibitors of the cell-wall. If our hypothesis is true, we reasoned that bedaquiline should not profile with cell-wall acting antibacterials when grown in media containing a fatty acid as its sole carbon source. We tested this hypothesis by comparing profiles of Mtb grown in standard rich growth medium or a growth medium with the fatty acid butyrate as the sole carbon source. We observed that morphological profiles are highly dependent on growth environment (Fig. S5A) with bedaquiline profiles resembling those from cell-wall acting antibacterials only when Mtb is grown in rich medium (Fig. S5B). These data support previous observations (6) that the mechanism of action of bedaquiline is dependent on metabolic state of Mtb.

### MorphEUS correctly classifies mechanisms of cellular death provoked by unknown drugs

Classification of morphological profiles using MorphEUS show that distinctive morphological patterns are induced in Mtb according to the terminal stress pathway, which may be the canonical pathway of action or proximal (downstream) effector, as in the case of moxifloxacin and bedaquiline. Because some downstream or off-target effects may be induced at high dose treatments (or likewise not overshadowed by other pathways at low dose), dose-dependencies may be another indicator of non-canonical effects. In support of this hypothesis, we observed strong dose-dependencies with morphological profiles of bedaquiline and moxifloxacin (Fig. S7).

Blinded to compound identity, we next used MorphEUS to identify pathways of action for three non-commercial antituberculars with known mechanisms of action. We mapped unknown compounds 1 and 2 as cell-wall acting; compounds 1 and 2 were nearest neighbors to ethionamide and ethambutol, respectively (Fig. 4). We unblinded the compound identities to compare their known mechanisms of action to those predicted by MorphEUS. These compounds (DG167 and its derivative JSF-3285) were validated through extensive biophysical, X-ray crystallographic, biochemical binding and spontaneous drug-resistant mutant studies to be inhibitors of cell-wall mycolate biosynthesis through specific engagement of the α-ketoacyl synthase KasA ((27), and Cell Chem. Biol, provisionally accepted). Taken together, our analysis of DG167 and JSF-3285 using MorphEUS has independently validated the target pathway of these two compounds and shown these analogs act through the same pathway of action.

**Figure 4.**
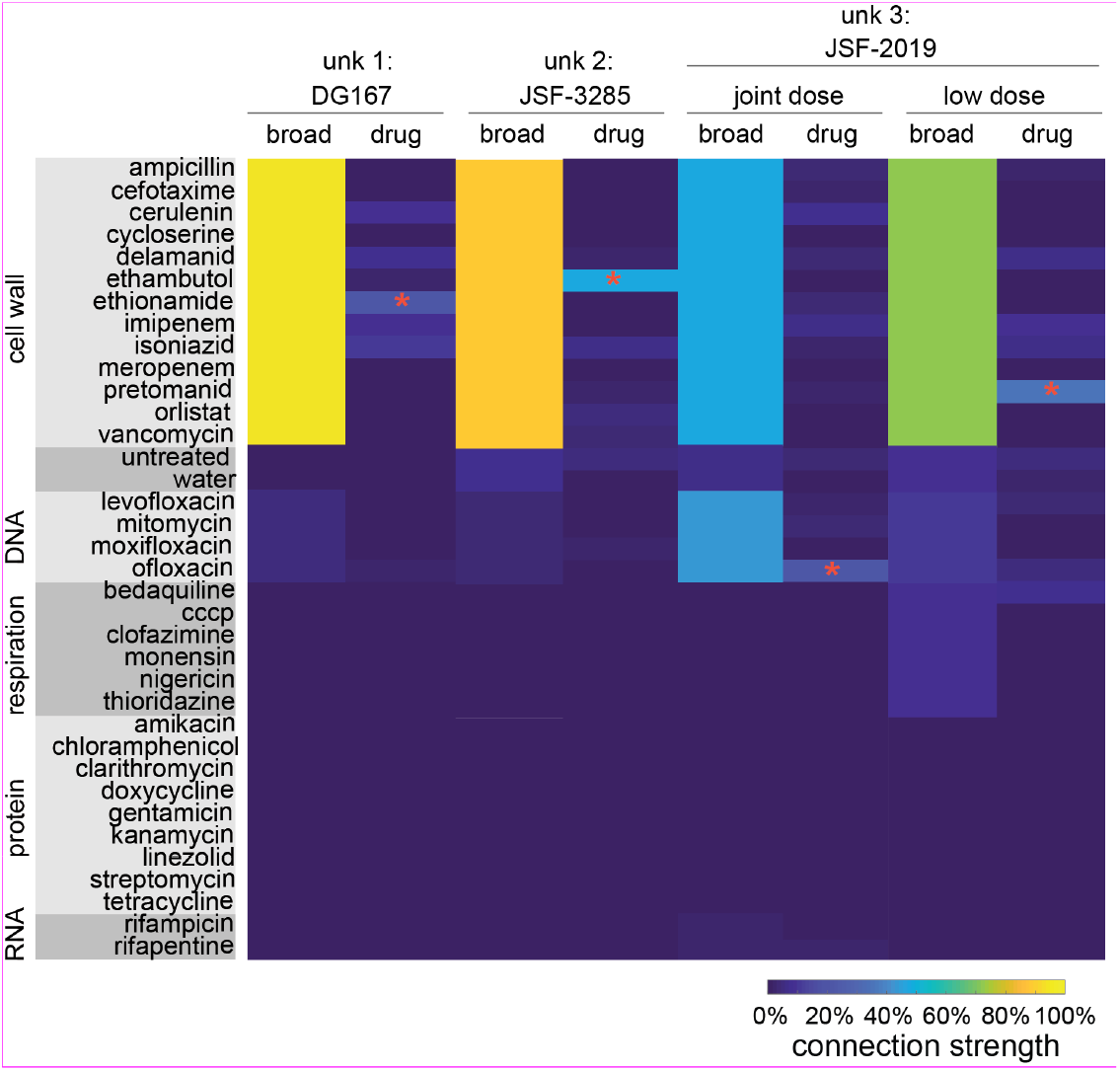
MorphEUS accurately predicts pathways of action of compounds when blinded to mechanism of action. cKNN profiles of broad drug categories and individual drugs for compounds with anti-TB activities (unk for unknown). Each column corresponds to a different compound or treatment dose (joint dose profile for DG167 and JSF-3825; n= 7300, 7160, 7742, 5150 from left to right). The most similar drug for each MorphEUS classification is indicated by the red asterisk.

The activity of the third unknown compound was harder to interpret. MorphEUS predicted unknown compound 3 to act on both the cell-wall and DNA and had ofloxacin as its nearest neighbor (via joint-dose profiles; Fig. 4). In contrast, MorphEUS analysis at low treatment dose mapped unknown compound 3 to cell-wall acting antibacterials with pretomanid as its nearest neighbor. Unknown compound 3 therefore displays dose dependent effects that become apparent in the joint profile suggesting that downstream off target effects are amplified upon increasing the treatment dose. We unblinded the compound to learn if our conclusions were corroborated with previous mechanistic studies performed. Unknown compound 3 was the recently published triazine JSF-2019 that resembles pretomanid in both its F420-dependent production of NO• and its ability to inhibit mycolic acid synthesis, albeit at a different step in the pathway (28). The mechanistic similarity of JSF-2019 with pretomanid validated the MorphEUS prediction of JSF-2019 acting like pretomanid at low dose but did not provide insight into the MorphEUS prediction of DNA targeting activity at high dose. We hypothesized that the production of NO• by JSF-2019 at high doses induces DNA damage through DNA alkylation (29). To test if JSF-2019 perturbs DNA processing pathways, we evaluated transcriptional profiles for ofloxacin- and JSF-2019-treated Mtb and found enrichment of co-regulated genes involved in DNA damage and repair (SOS response and purine synthesis), demonstrating Mtb experiences DNA damage when treated with JSF-2019 (Fig. S8). A detailed analysis of JSF-2019 resistant mutants (28) uncovered the presence of mutations in *rv2983* and *rv2623* (30–32). Mutations in *rv2983* have previously been found to generate resistance to fluoroquinolones (30) while overexpression of Rv2623 has been linked to exposure of Mtb to ofloxacin or moxifloxacin (31, 32). Together these analyses show that JSF-2019 has dual, dose-dependent pathways of action where JSF-2019 damages the cell-wall at low doses and DNA at high doses. We conclude that MorphEUS has enabled us to efficiently focus the analysis of resistance and transcriptomic data to define critical downstream effects that contribute to the mechanism of JSF-2019.

## Discussion

Unlike other bacterial species such as *E. coli*, *B. subtilis*, and *M. smegmatis,* the morphological response of Mtb to drug treatment was subtle and confounded by high levels of heterogeneity in morphological features. We do not understand why shifts in morphology are so mild in Mtb but hypothesize that the slow growth of Mtb may diminish the readily observable morphological response to antibacterial compounds i.e. that catastrophic cellular stress may proceed slowly and less chaotically. We designed MorphEUS, a new morphological profiling pipeline, to overcome these challenges and enable us to profile drug action by cellular damage as manifested in physical changes such as cell shape, permeability, and organization. The subtle changes in features were drug specific, dose dependent, and influenced the heterogeneity in Mtb’s morphological response (Fig. 1B, Fig. S4), providing us with a valuable tool in which to increase the information provided by cellular damage as a marker of drug action. MorphEUS therefore incorporates cell-to-cell heterogeneity in morphological features as key measurements to distinguish among affected pathways of action. In addition, MorphEUS accounts for subtlety among features with multiple classification trials of k-nearest neighbor mapping to preserve higher-dimensionality in classification.

In applying MorphEUS to a set of 34 known antibiotics and three blinded, non-commercial antibacterials, we found that MorphEUS grouped antibacterials by their killing activity, which may be the primary target pathway or off-target effects. Most of the antibacterials profiled with their known pathway of action, but we identified three drugs (moxifloxacin, bedaquiline, and JSF-2019) where killing activity mapped instead to off-target effects. Bedaquiline and moxifloxacin are known as inhibitors of respiration and DNA synthesis, respectively, yet both clustered with cell-wall-inhibiting drugs. For bedaquiline, we found that the cell-wall damage effect was specific to metabolic state and was a secondary effect for cells carrying out glycolysis and was not observed for cells utilizing fatty acids as a carbon source. We corroborated the apparent cell-wall acting activity of moxifloxacin at high doses using transcriptional analysis. There is increasing evidence that poly pharmacologies and off-target effects significantly contribute to the bactericidal activity of a drug (4, 20–22). For example, recent reports in *M. absessus* and *M. bovis* have shown that treatment with cell-wall acting compounds lead to toxic intracellular accumulations of ATP - a downstream effect that is independent of the cell-wall activity of the drug (21, 22). Another example of the bactericidal activity from an off-target effect can be found in *E. coli* and its production of toxic free radicals following treatment with inhibitors of protein, DNA, and cell-wall synthesis (4). We anticipate that expansion of MorphEUS analyses to larger drug sets and more metabolic states will identify more important off-target effects.

MorphEUS profiling, like all cytological profiling techniques, is data-driven and based on classification among a pool of other profiles. Therefore, MorphEUS is sensitive to both the breadth and depth of the antibacterials used to create the profiles. Nonetheless, MorphEUS is a powerful tool to rapidly generate hypotheses about the pathways of action for antibacterials, compounds in development and drugs in specific growth environments. These profiles and hypotheses are based on morphological rather than molecular signatures, and well complement transcriptomic and genetic approaches by focusing how to evaluate these large systematic datasets. Through the lens of off-target effects from MorphEUS, we examined transcriptional profiles for treatment with moxifloxacin and JSF-2019; these transcriptional data support the MorphEUS-generated hypotheses that these compounds damage bacilli through off-target effects and not only effects associated with the primary target of these compounds. These findings highlight the ability of MorphEUS to link the morphological response of Mtb to the proximal cause of death in Mtb through interpretable hypothesis generation.

We have shown that the MorphEUS pipeline identifies drug pathways of action, also revealing off-target and downstream drug effects that are proximal to antibacterial action. We anticipate that the systematic application of MorphEUS will further reveal drug polypharmacologies and detail how the killing activity evolves as one proceeds from drug engagement of cellular target(s) to cell death. With a well-sampled dataset of treatment profiles to benchmark drug-target pathways, MorphEUS can be applied to rapidly identify new compounds that have unique mechanisms of action, thereby accelerating the drug development pipeline for tuberculosis. We expect that the success of MorphEUS in profiling drug action in an organism with significant inherent heterogeneity and subtle cytological responsiveness is an indicator of the analysis’s translatability to other pathogens and cell types.

## Materials and Methods

### Bacterial strains

*Mycobacterium tuberculosis* strain used in this study was Erdman. *Mycobacterium smegmatis* strain used in this study was derived from mc^2^155. *Escherichia coli* strains used in this study were derived from DH5α.

### Growth conditions

*M. tuberculosis* cells were cultured in 7H9-rich medium consisting of 7H9 broth (ThermoFisher, DF0713-17-9) with 0.05% Tween-80 (ThermoFisher, BP338-500), 0.2% glycerol (ThermoFisher, G33-1), and 10% Middlebrook OADC (ThermoFisher, B12351). Frozen 1ml stocks were added to 10 ml 7H9-rich medium and grown with mild agitation in a 37°C incubator until the culture reached an OD_600_ of ~0.4-0.7. The bacteria were then subcultured into 10 ml of fresh medium to an OD_600_ of 0.05 and grown to an OD_600_ of ~0.4-0.7. At this time the cells were plated onto 96-well plates containing drugs at the predetermined amounts (see below). Drug-treated plates were incubated at 37°C in humidified bags until fixation.

*M. tuberculosis* cells were adapted to low pH medium by first growing and subculturing the cells once in 7H9-rich medium (as described above) followed by centrifugation and resuspension in 7H9-rich medium supplemented with 100 mM MES hydrate (SigmaAldrich, M2933) HCL adjusted to pH of 5.8. Cells were subcultured once in 7H9-rich-low pH medium before plating.

*M. tuberculosis* cells grown with butyrate or cholesterol as their sole carbon source were cultured in 7H9-base medium (7H9 broth with 0.05% Tyloxapol, 0.5 g/L Fatty Acid-free BSA, 100 mM NaCl, 100 mM MOPS buffer (SigmaAldrich, M3183), and HCL adjusted to pH 7.0) supplemented with either 10 mM sodium butyrate (SigmaAldrich, 303410) or 0.2 mM cholesterol (SigmaAldrich, C8667). Sodium butyrate was added directly to the 7H9-base medium while cholesterol was dissolved in a 50/50 (v/v) mixture of tyloxapol and ethanol to obtain a 100 mM stock solution as previously described (23). Bacteria grown in butyrate medium were grown and subcultured once in 7H9-rich medium before centrifugation and resuspension in butyrate medium. The cells were subcultured once using fresh butyrate medium before they were aliquoted into tubes (1 ml each) which were stored at −80°C until use. Frozen stocks were started and subcultured in butyrate medium before plating. Bacteria grown in cholesterol medium were grown and subcultured once in 7H9-rich medium before centrifugation and resuspension in cholesterol medium to an OD_600_ of ~0.2. The bacteria were plated upon reaching an OD_600_ of ~0.4. *M. smegmatis* cells were cultured in 7H9-rich medium supplemented with Middlebrook ADC (ThermoFisher, B12352). 100 μl frozen stocks were added to 10 ml of 7H9-rich-ADC medium and subcultured once before use. *E. coli* cells harboring plasmids used in this study were grown in LB broth containing appropriate antibiotics (50 μg/ml hygromycin or 25 μg/ml kanamycin).

### Drug treatments

For time-dose-response profiling, drugs were loaded into 96-well plates with the HP D300e digital drug dispenser. Each drug used in the study was reconstituted, depending on drug solubility, in water, DMSO, 1N NaOH, or methanol solubility at a concentration between 2.5 and 100 mg/ml (Table S1). Reconstituted drugs were then aliquoted in single-use sterile tubes and stored at −20°C until use. The percentage of DMSO for all drug treatments was between 0.00045-0.75% except for ampicillin, tetracycline, chloramphenicol, and thioridazine high dose treatments, where DMSO percentage did not exceed 1.5%. Even at 1.5% DMSO, profiles of untreated cells and DMSO-treated cells were indistinguishable. To determine whether solvents elicited morphological changes that would impact the profiling, we tested whether cells treated with each control condition (DMSO at different concentrations, and the highest levels used in the other solvents: 0.3% 1N NaOH, 0.1% methanol, or 3% water) profiles with each other and untreated samples, which would suggest that the solvents were not drivers of morphological changes. We selected a feature set based on the high dose MorphEUS analysis, keeping features that were used in over half of the classification trials, resulting in a set of 28 features. Using the untreated profiles from the high dose analysis, we performed TVN normalization on the range of DMSO treatments followed by PCA. We searched for the first nearest neighbor for each individual treatment to see if the same treatment groups were nearest neighbors with each other. We found no likeness (e.g. 100% confusion) between the same treatment groups (e.g. that the controls were not identifiable into similar treatment groups), suggesting that solvents alone did not induce strong morphological effects.

### Inhibitory Concentration 90 (IC90) determination

*M. tuberculosis* and *M. smegmatis* cultures were grown from frozen aliquots added to their respective 7H9-rich medium, and subcultured once as described above. Once grown to an OD_600_ of ~0.4-0.7, the cells were diluted to an OD_600_ of 0.05 and added to 96-well plates containing drugs in a twofold dilution series for 9 concentrations. Each treatment series contained an untreated well as a control. All IC90 determinations were performed in biological triplicate. To avoid plate effects, wells around the perimeter of the plate were not used. An initial OD_600_ plate read was performed immediately for each *M. smegmatis* plate, while for *M. tuberculosis* cultures this was performed after allowing the bacterial cells to settle overnight. A second plate read was performed for *M. smegmatis* after 24 hours and *M. tuberculosis* after five days of incubation. Growth inhibition curves were generated by subtracting the initial reads from the final reads and then normalizing the data to untreated controls. The 90% growth inhibitory concentration (IC90) was defined as the drug concentration that inhibited at least 90% of all bacterial growth.

### Fixation of antibiotic treated Mtb-bacilli

After the designated treatment times (overnight unless otherwise noted), Mtb cultures were fixed in paraformaldehyde (Alfa Aesar, 43368) at a final concentration of 4% and transferred to clean 96-well plates. The plate was surface decontaminated with vesphene IISE (Fisher scientific, 1441511) and sealed with Microseal ‘F’ foil seals (Biorad, MSF1001). The duration of fixation was one-hour total. After fixation, the cells were washed twice with 100μl of PBS (ThermoFisher, 20012-027) + 0.2% Tween-80 (PBST), then resuspended in 100 μl of PBST, sealed (ThermoFisher optically clear plate seals, AB1170) and stored at 4°C until staining and imaging.

### Staining and fluorescent imaging of Mtb cells

All staining was performed in 96-well plates with 50μl of fixed Mtb cells diluted in 50 μl of PBST. Staining was performed with 0.6 μg of FM4-64FX (ThermoFisher, F34653) and 15 μl of a 0.1 μM SYTO 24 (ThermoFisher, S7559) stock in each well containing PBST and fixed bacilli. The plate was then incubated at room temperature in the dark for 30 minutes. Once stained, the cells were washed once with an equal volume of PBST and resuspended in 30 μl of PBST. Stained Mtb were spotted onto agar pads (1% w/v agarose; SigmaAldrich A3643-25G). Images were captured with a widefield DeltaVision PersonalDV (Applied Precisions) microscope. Bacteria were illuminated using an InsightSSI Solid State Illumination system with transmitted light for phase contrast microscopy. SYTO 24 was imaged using Ex. 475nm and Em. 525nm. FM4-64-FX was imaged with Ex. 475nm and Em. 679nm. Montage images were generated using a custom macro that captures 25 individual fields of view per image. Two technical replicate images were taken from each sample for a total of 50 images per biological replicate. Three biological replicates were generated for each drug treatment. Images were recorded with a DV Elite CMOS camera for all three channels.

### Generation of RpoB-GFP strain

A strain of *rpoB-gfp* in the *M. smegmatis* mc^2^155 background was generated using the ORBIT recombineering system developed by Murphey et al. 2018 (24). Briefly, a frozen aliquot of *M. smegmatis* was grown and subcultured once as described above. Upon reaching mid-log phase the cells were washed twice with 10% glycerol (Fisher Scientific, G33-1) and electroporated with pKM444. The plasmid pKM444 allows for ATC inducible expression of Che9c RecT and Bxb1 integrase phage proteins and harbors a kanamycin resistance cassette. Transformants were selected for on Middlebrook 7H10 plates (ThermoFisher, BD 2627) with ADC containing 25 μg/ml of kanamycin (VWR, 0408-10G). A control without plasmid was also plated to ensure proper kanamycin selection. The pKM444 harboring strain of *M. smegmatis* was then grown to an OD_600_ of 0.5 in 7H9-rich-ADC medium containing 25 μg/ml of kanamycin. Once the desired OD_600_ was reached, anhydrotetracycline (ATC; Fisher Scientific, 13803-65-1) was added and the cells were incubated with gentle agitation until an OD_600_ of 0.8 was reached. The cells were then washed with glycerol as described above and electroporated with 1μg of an *rpoB* targeting oligo harboring an *attP* sequence (see below) and 0.2 μg of the non-replicating *GFP* harboring plasmid pKM468. pKM468 contains an *attB* recombination downstream of the *egfp* gene for N-terminal translational fusions, lacks a mycobacterial origin of replication and harbors a hygromycin resistance cassette. -Oligo+plasmid and -oligo-plasmid controls were also performed as negative controls. Transformations were recovered in 1 ml of 7H9-rich-ADC medium, incubated for three hours then plated on 7H10-ADC plates containing hygromycin B at 50 μg/ml. The presence of the C-terminal GFP translational fusion to RpoB was validated by fluorescence microscopy using the FITC (Ex. 475nm Em. 525nm) channel as described above. The *rpoB* targeting oligo sequence was: 5’GCACGTAACTCCCTTTCCCCTTGCGGGTGTTGAAACTTGACTACTGAGGCGGTCTTCGGACGAGGCTCTAGGTTTGTACCGTACACCACTGAGACCGCGGTGGTTGACCAGACAAACCCGCGAGATCCTCGACGGACGCGGATTCGTTGCGCGACAGGTTGATTCCCAGGTTCGCGGCAGCGCGCTCC 3’.

### Live-cell microscopy

*M. smegmatis* cells expressing RpoB-GFP were grown overnight from frozen 100μl aliquots in 10 ml of fresh 7H9+ADC. The bacteria were subcultured once and allowed to reach mid log phase (OD_600_ ~0.5-0.7). The culture was then filtered to remove aggregates of bacteria and loaded into a custom polydimethylsiloxane (PDMS) microfluidic device as previously described (25). Fresh medium was delivered to cells using a microfluidic syringe pump. The microfluidics device was attached to a custom PDMS mixing device for delivery of drug for the duration of time described below and then placed on an automated microscope stage inside an environmental chamber that was maintained at 37°C. The bacteria were imaged for a total of 26 hours using a widefield DeltaVision PersonalDV (Applied Precision, Inc.) with a hardware-based autofocus. Antibacterial compounds were introduced to the *M. smegmatis* after a 10-hour growth phase. Drug treatment lasted for 6 hours and was followed by a 10-hour recovery phase.

### Transcriptional profile analysis

The transcriptional profile of JSF-2019, bedaquiline and moxifloxacin were obtained from GSE126718 (26), GSE43749 (6) and GSE71200 (10), respectively. The transcriptional profiles of other compounds were extracted from GSE1642 (11). The 166 genes which were significantly co-upregulated and co-downregulated by both JSF-2019 and ofloxacin (judged by fold change >1.5 and FDR p-value <0.01) were selected from the transcriptional profile. *rv0560c*, which encodes a benzoquinone methyltransferase involved in xenobiotics metabolic detoxification (27, 28), was removed from the list due to its extremely high induction fold in JSF-2019 rather than in other compound treatments. A hierarchical clustering analysis was applied to the 165 genes according to Euclidean distances via R packages pheatmap and ggplot. The function and pathway enrichment analysis of the 155 genes were performed via gene ontology resource (http://geneontology.org/).

## MorphEUS analysis pipeline

### Overview of Mtb morphological profiling analysis (MorphEUS)

The MorphEUS analysis pipeline (Fig. 2) is as follows:

#### Feature quantification

1) Single cell measurements extracted from MicrobeJ are imported to MATLAB to 2) undergo quality control before bulk analysis. 3) Variables describing the cell-to-cell variation within each feature are calculated prior to feature selection and profile classification.

#### Classification trial

4) Non-redundant features are iteratively selected. 5) Normalization is performed across replicates to decrease experimental noise (TVN). 6) PCA is performed on the batch-normalized, reduced feature dataset and followed by 7) KNN analysis on the PCA scores matrix. KNN analysis classifies drugs (e.g. treatment groups) by identifying the nearest neighbors (most similar profiles). These nearest neighbors are represented as a matrix or a network diagram (graph).

##### Classification consensus [consensus KNN (cKNN)]

To overcome the fragility of feature selection, we generate a consensus of multiple (70) classification trials, wherein each trial utilizes a different, stochastically selected set of 80 untreated samples for feature selection and batch normalization. The resulting cKNN may be visualized using a network diagram where edges between drugs are color-coded according to how often their profiles were nearest neighbors (% of total trials) or a matrix that describes the frequency of drug-drug links among trials (Fig. 3). The most similar drug profiles are linked in a large number of the trials. The connectivity maps and corresponding summary heatmaps are used to make informed predictions about the target pathway of an antibiotic.

#### Image segmentation and feature extraction

Before image segmentation, we used the ImageJ plugin BaSiC to ensure an even distribution of illumination in all channels across the image (29). Image segmentation was performed using the ImageJ plugin MicrobeJ (v 5.13l (1)), extracting seven features from phase contrast and nine from each fluorescent channel using custom settings, resulting in a total of 25 features (Table S2) (30). The image segmentation in MicrobeJ is computationally demanding and therefore was run on a high-performance computing cluster.

#### Blur thresholding

All data were organized and analyzed using custom scripts in MATLAB (2019a). Out-of-focus bacilli were identified from the transverse phase-contrast profile of each bacterium and discarded. The profile of an in-focus, well-segmented bacillus has a gaussian distribution with high intensity around the edge of the bacterium, followed by a steep drop after the edge. Blurry cells were filtered out with the following criteria: goodness of fit to a gaussian distribution, if the minimum point was off center, unexpected local maxima, difference in intensity between minimum and maximum values, difference in intensity between edges, and slope of the edges. With exception to the fluorescent foci counts, the median, first quartile, third quartile, and interquartile range (IQR) were calculated for each feature to account for the population distribution, resulting in 94 features total. Distribution features were not calculated for foci count features because these measurements are discrete, not continuous, features. Features were then normalized, dividing by the largest value for the feature across all treatments.

#### Typical Variance Normalization

Typical Variation Normalization (TVN) aligns the covariance matrices produced by PCA of untreated control data from each experimental plate, or batch, and applies this transformation batch-by-batch to allow for less biased comparison of the drug-treated cells across plates and replicates (31). An abbreviated version of TVN was applied to reduce batch effects from imaging. First PCA is performed on the untreated controls. Each axis is scaled to have a mean of zero with variance of 1. This transformation is then applied to the entire dataset, including treated and untreated samples (Fig. S3). To perform TVN, we dedicated 25% of our samples in every experiment and imaging session to be untreated controls. Each classification trial begins with stochastically selecting 80 untreated controls; thus, feature sets were restricted to a maximum of 79 features since a PCA transform with n samples can only have n-1 features.

#### Principal Component Analysis

PCA was performed on the normalized data (feature selected or not, as indicated) using the built-in MATLAB function. In the BCP pipeline, PCA reduces the dimensionality of the data, allowing variance across all of the features to be visualized in 3 or fewer dimensions. After accounting for heterogeneity with batch normalization and including features of variation into the profiles, some drug clustered observed, especially among cell-wall acting antibacterials (Fig. S1A lower). By PCA, bedaquiline clustered with cell-wall acting drugs in standard, rich growth medium (Fig. S1A lower and Fig. S5B left) but not in fatty acid-rich growth medium (Fig. S5B right).

#### K-nearest neighbors

K-nearest neighbors (KNN) analysis was implemented using the cosine distance metric and the *knnsearch* MATLAB function. K was set to 1, thus only the first nearest neighbor was identified. For our setup, we took the median PCA score from the three replicates for each drug as inputs for the KNN analysis. The KNN algorithm finds the k-nearest neighboring points where the cosine distance between PCA scores is shortest. MATLAB defines the cosine distance as one minus the cosine of the included angle between points. We observed that feature selection was dependent on which untreated samples were included in the TVN batch-to-batch normalization process (80 from 117), suggesting there are many good solutions, or feature sets that can lead to similar profiling of the drug target. To ensure our classification was not overfitting the data depending on which untreated samples were included in the analysis, we took a stochastic approach. We define the application of PCA and then KNN analysis on a particular set of reduced features as a classification trial. The MorphEUS pipeline steps 4-7 were repeated for 70 classification trials, each including a different randomly selected set of 80 untreated controls (classifications converged by 70 trials; Fig. S8).

#### Iterative feature selection

To reduce overfitting and noise in our 94 variable feature set, we utilized the Minimum Redundancy Maximum Relevance (mRMR) feature selection algorithm (32). Here we customized previously published MATLAB code to perform mRMR feature selection using the mutual information difference scheme (32). Because the algorithm rank orders variables and does not automate the selection of the optimal number of features, we implemented an iterative feature selection method that rewards runs that result in more drug-drug connections with same target pathways (Table S1). Since we begin with 94 features but are limited to 79 variables by our TVN analysis, mRMR was used to rank order the top 79 features. Starting with the 79 rank-ordered features, we removed each feature individually and performed TVN, PCA and KNN analysis on the remaining feature set. Success of the feature set was quantified by accuracy of the KNN in linking drugs belonging to the same broad category assigned by literature review (see Table S1). The feature set that resulted in greatest model accuracy was selected, and the variable removal process was repeated until maximal prediction performance was reached. On average, these iteratively determined feature sets contained 38 variables.

#### Consensus KNN (cKNN)

The cKNN results compiled from all 70 classification trials were visualized using a network map and heatmap, where edge color and grid square color, respectively, corresponds to how frequently two drug profiles were identified as nearest neighbors. Because our goal was to identify similar treatment profiles, connections between profiles in the cKNN were made undirected and the drug-drug categorization matrices (such as Fig. 3B left) are symmetric. These visuals allow for classification of the target biological pathway of each drug based on the robustness of phenotypic similarities between drug profiles can easily be evaluated. All maps and plots were generated in MATLAB (2019a).

#### Comparison to random

As a comparison to model accuracy by random change, we tested how accurate MorphEUS was when the drug categories were randomly assigned. To do so, the labels for the drugs in the final cKNN were randomly swapped, resulting in 22% accuracy compared to 94% for the joint dose MorphEUS analysis.

#### Cross validation and classification of unknown compounds

To test the strength of our model, we performed cross validation. This was done by removing one of the drugs out of our 34-drug set and running the remaining 33 drugs through the MorphEUS pipeline. The PCA transformation created by the 33 drugs was applied to the TVN-normalized, removed drug and KNN analysis was performed. At the end of the 70 trials, a cKNN was created and the pathway of action of the cross validated drug was classified in accordance to its strongest drug connections and their corresponding pathway(s) of action as classified in Table S1.

#### Low dose, high dose, and joint dose profiles

Mtb cytological features are dependent on drug target but also treatment dose and duration (Fig. S6). This raised the possibility that morphological profiles from a low dose of treatment or a joint profile of low (0.25xIC90) and high (3xIC90) dose treatments would improve the accuracy of drug classification using the full drug set and subsequent cross validation. To investigate whether a joint dose profile best describes the variation in the morphological response in Mtb, the full 94 feature datasets from both drug doses were concatenated, resulting in 188 total features. We also applied MorphEUS to low dose and high dose treatments as separate profiles. We observed high accuracy using each of the dose treatments (low, high, and joint as 97, 91, and 94% respectively), but the joint dose profiles were better cross validated (76%) compared to high (68%) and low (62%) dose MorphEUS. We therefore use joint dose profiling as the default for MorphEUS.

#### Classification of unknown compounds

We apply new compounds to MorphEUS in the same manner as cross validation, only the MorphEUS pipeline is done on the full 34 drug set and the unknown is added to the set for the final KNN during each classification trial.

#### Statistical Analysis

We performed the Kruskal-Wallis test to identify drug treatments that induce significantly different morphological features compared to untreated cells in rich medium (Fig. 1B and S5A). While the MorphEUS pipeline utilizes population-based features, the Kruskal-Wallis test was applied to the features of individual cells (n=1625-3983 for rich, n=1029-6733 for conditions). The Kruskal-Wallis test was applied to each drug and/or environmental condition individually, per feature. In each case the null hypothesis was that the median feature value for Mtb exposed to a specific drug and/or environmental stress was drawn from the same distribution as the median feature value for the untreated controls.

## Acknowledgments

We thank R. Abramovitch, J. Seeliger, D. Warner, V. Dartois, and S. Tan for insightful discussion and B. Kana, K. Rhee, and C. Stallings for valuable discussion and critical reading of the manuscript. The plasmids pKM444 and pKM468-EGFP were gifts from Kenan Murphy and Chris Sassetti (Addgene plasmid # 108319; http://n2t/net/addgene:108319; RRID:Addgene_108319 and Addgene plasmid # 108434; http://n2t.net/addgene: 108434; RRID:Addgene_108434 respectively)

## Author Contributions

T.C.S., K.M.P., X.W., J.S.F, and B.B.A. designed the study. T.C.S., K.M.P., and M.E.M conducted the experiments. T.C.S., K.M.P., M.C.O., X.W., J.S.F., I.R., S.H., D.M.A., and B.B.A. designed and implemented the analysis. T.C.S., K.M.P., M.C.O., M.E.M. X.W., J.S.F, and B.B.A wrote the paper, which was edited by all authors.

