## Supplementary figures for "Morphological profiling of tubercule bacilli identifies drug pathways of action"

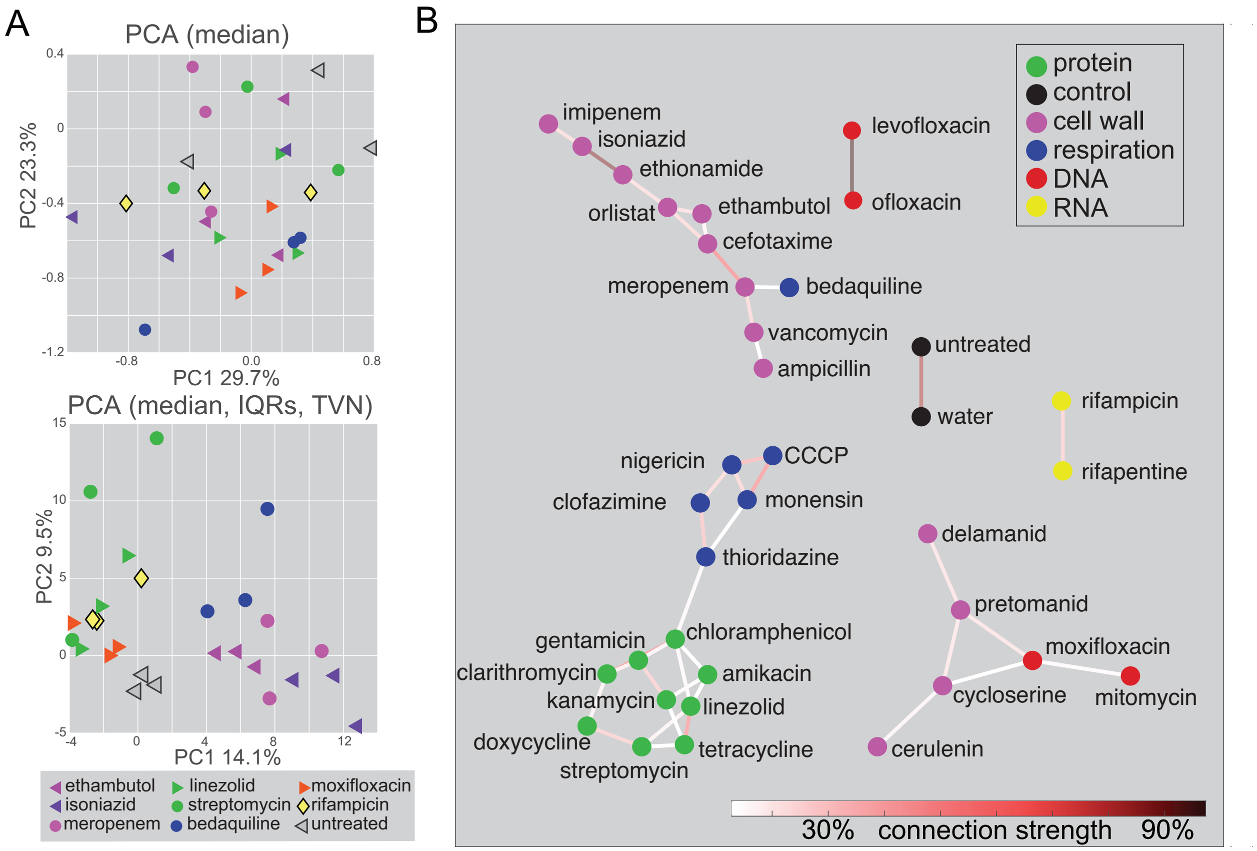


**Figure S1. Inherent heterogeneity of Mtb and subtle morphological features requires multistep, multivariable analysis to define cytological profiles.** (A) Principal component analysis of seven drug treatments resulting from established BCP analysis methods using feature medians alone (1-3) (top) and after incorporating metrics of heterogeneity [quartile 1, quartile 3, and interquartile range (IQR)], feature selection, and batch normalization (TVN; bottom). n=3032-9651 for each treatment group over biological triplicate. TVN aligns the covariance matrices resulting from PCA and whitening of untreated control data from each batch and applies this transformation batch-by-batch to enable comparison across plates and replicates. (B) Consensus k-nearest neighbor (cKNN) connectivity map for high dose drug treatment of Mtb. Drug nodes are colored by broad target category and links are color-coded by how frequently profiles are nearest neighbors in the classification trials (connections are shown for links >12%).

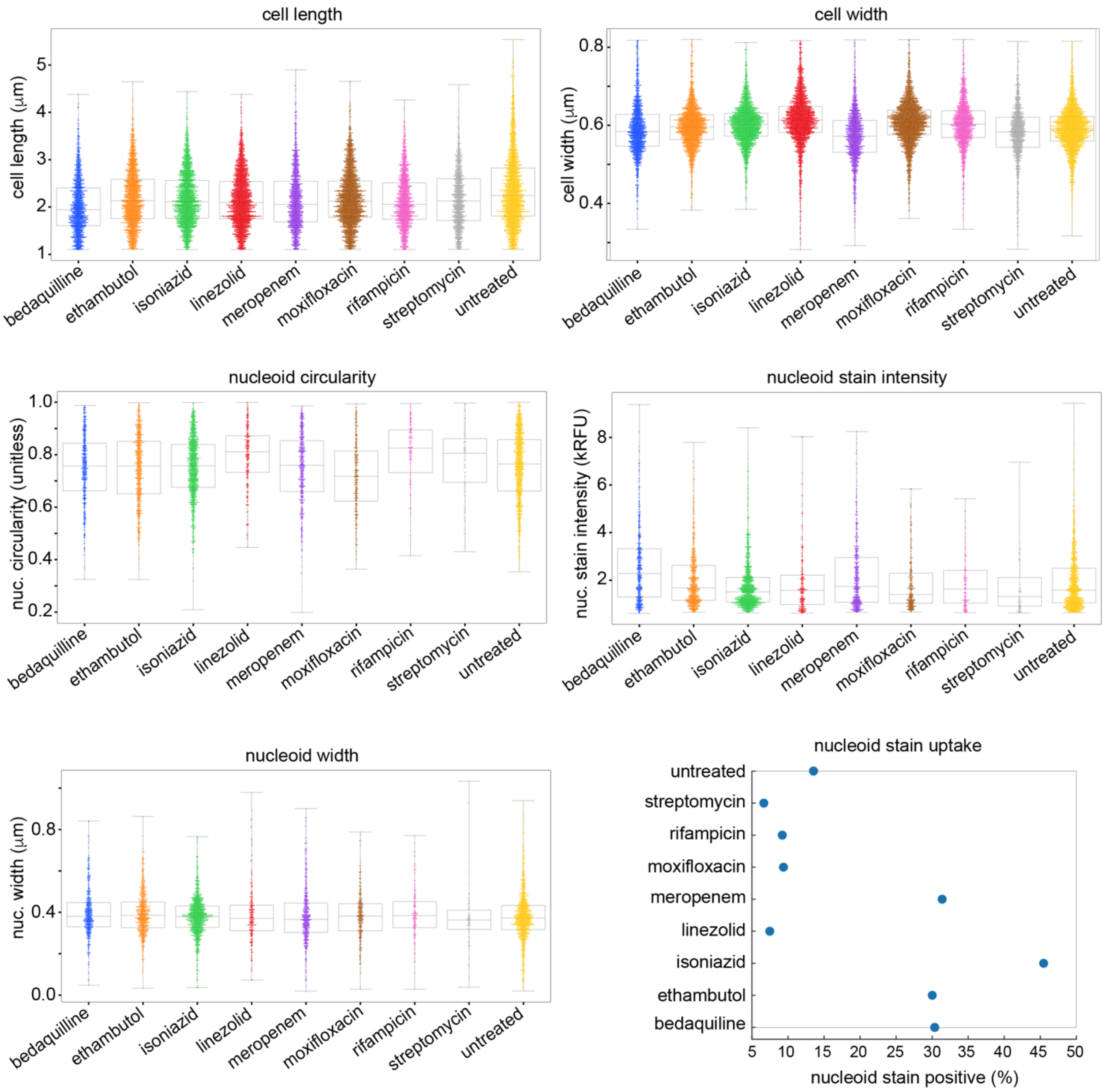


**Figure S2. Innate cell-to-cell heterogeneity in Mtb morphologies.** Dot and box plots of Mtb morphological features in untreated (yellow) and drug-treated bacilli. Many bacilli are not stained with STYO24, therefore only non-zero values are shown for nucleoid features. Cellular heterogeneity is apparent among the most basic cytological features, including cell length, intensity of nucleoid staining, and nucleoid width, even in untreated Mtb (CV=0.31, 0.64, 0.32, respectively). All violin plots were generated using the statistical data visualization library Seaborn for Python.

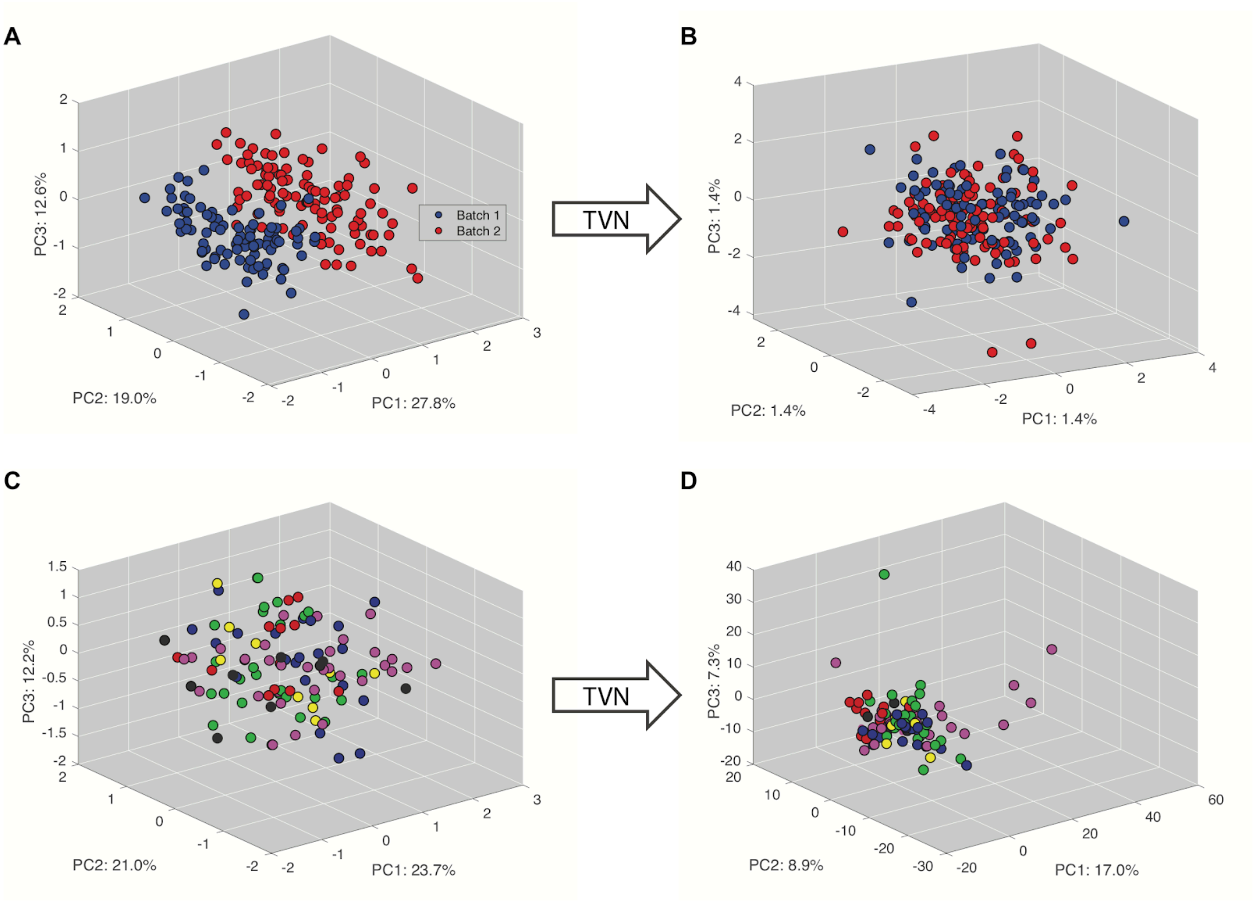


**Figure S3. Correction of batch-to-batch heterogeneity.** Principal component analysis (PCA) of untreated data labeled by batch. (A) PCA prior to TVN normalization. (B) PCA after TVN normalization. PCA of all 34 drug treatments labeled by target pathway using all 94 features. (C) PCA prior to TVN normalization. (D) PCA after TVN normalization. No feature selection was applied.

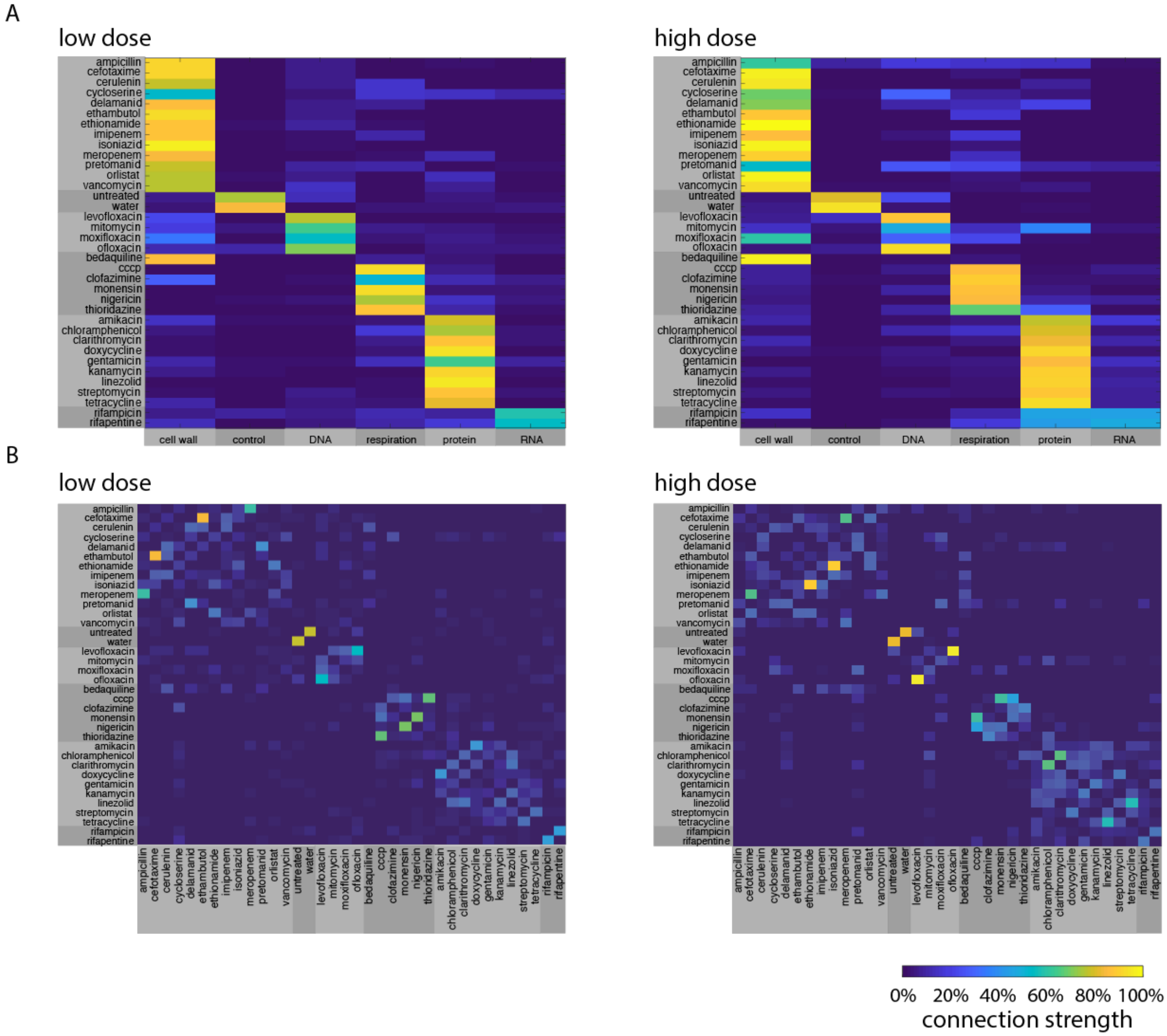


**Figure S4. Categorization of drug profiles using MorphEUS.** (A) Broad categorization for low dose (Left) and high dose (Right) profiles are represented as heatmaps. The connection strength is determined by the strength of the consensus k-nearest neighbor analysis. 100% strength indicates that the nearest neighbor identified in every iteration was from the same target category while 0% indicates that the drug of interest was never a neighbor to a drug in that pathway. Drugs are listed in the rows and collection of drugs within the broad target pathways are listed in the columns. (B) Heatmaps of drug nearest neighbor paring for low dose (left) and high dose (right) treatment. The heatmaps are described in Fig. 3. The high dose profiles were 91% accurate for assignment of broad drug category (compared to 26% for randomized categorization) with 68% accurate cross validation. The low dose profiles were 97% accurate for assignment of broad drug category with 62% accurate cross validation.

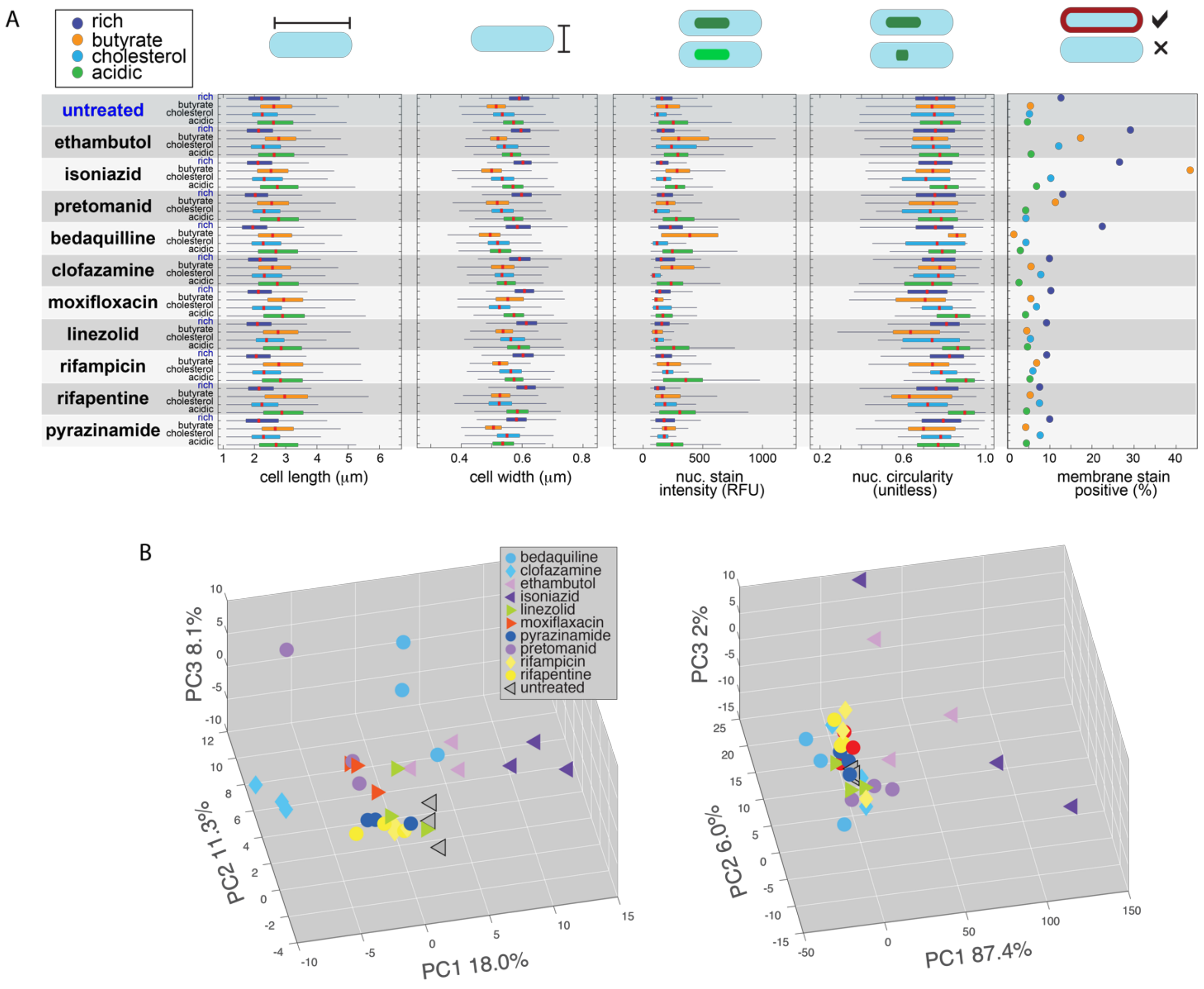


**Figure S5. Morphological features and response to drug treatment are dependent on growth conditions.** (A) Mtb were treated with each antibiotic (or untreated control) following adaptation to different growth media: rich (blue), butyrate as the sole carbon source (orange), cholesterol as the sole carbon source (cyan), and low pH in rich medium (green) as described in Methods. Red lines mark the medians, thick whiskers mark the 25-75th percentiles, and the thin whiskers extend the range of parameters that are not outliers (n=1029-6733). (B) PCA of morphological profiles in Mtb treated with antibiotics. Mtb cells were grown in media containing carbohydrates (left) or butyrate (right) as the sole carbon source. Bedaquiline (light blue circles) clusters close to the cell wall acting antibiotics (purple triangles) in media containing carbohydrates (left) but not in butyrate (right).

A

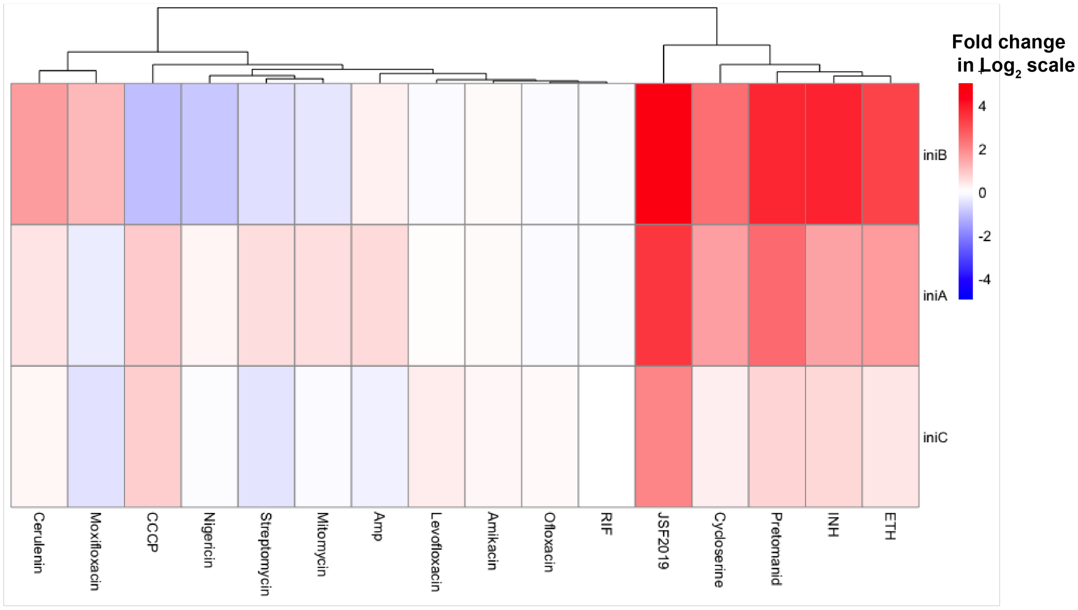


B

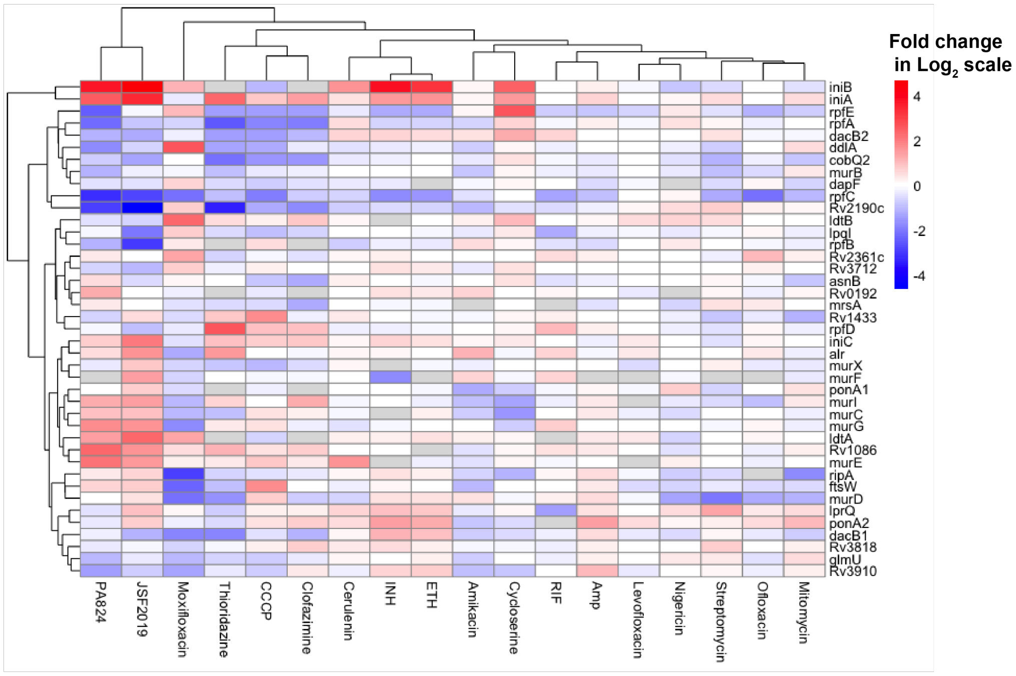


**Figure S6. Transcriptional response of Mtb to antibacterials in genes involved in cell wall damage and biosynthesis.** A) Expression profiles of the *iniBAC* operon in response to 16 antibacterials. Known inducers of the *iniBAC* operon isoniazid (INH), ethambutol (ETH), and pretomanid are plotted as controls B) Expression profiles of genes involved in peptidoglycan biosynthesis in response to treatment with antibacterials.

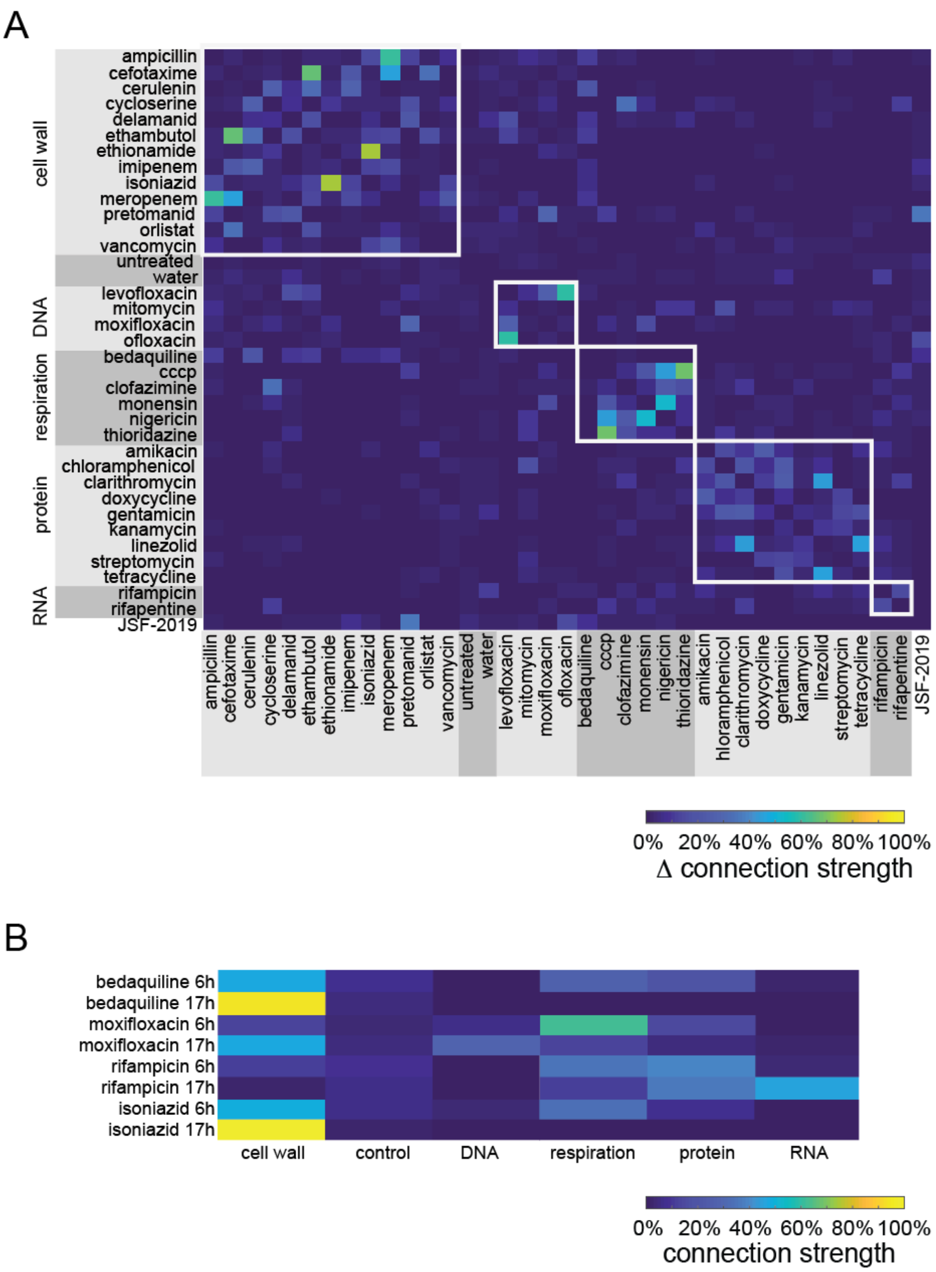


**Figure S7. Time and dose-dependency of profiles.** (A) Drug-drug matrix of the absolute difference in connection strengths as determined by MorphEUS analysis between low and high dose profiles with JSF-2019 applied. (B) Broad categorization of bedaquiline, moxifloxacin, rifampicin, and isoniazid at 6h mapped onto the joint dose profiles at ~1 doubling time (Fig. 3). The 17h profiles are shown as a point of comparison.

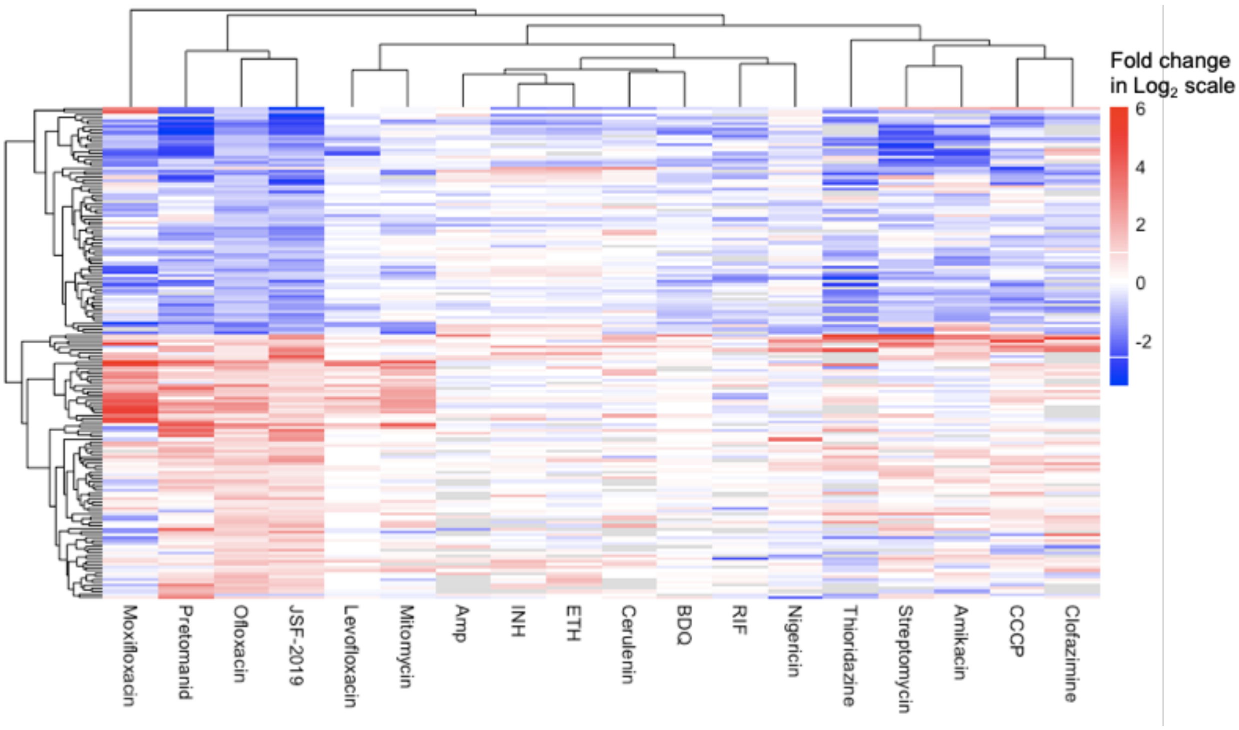


**Fig. S8. Transcriptional analysis of 165 genes co-regulated by JSF-2019 and ofloxacin.** The gene expression profiles of 165 genes that were significantly co-regulated by JSF-2019 and ofloxacin by >1.5-fold were analyzed via hierarchical clustering analysis by Euclidean distance.

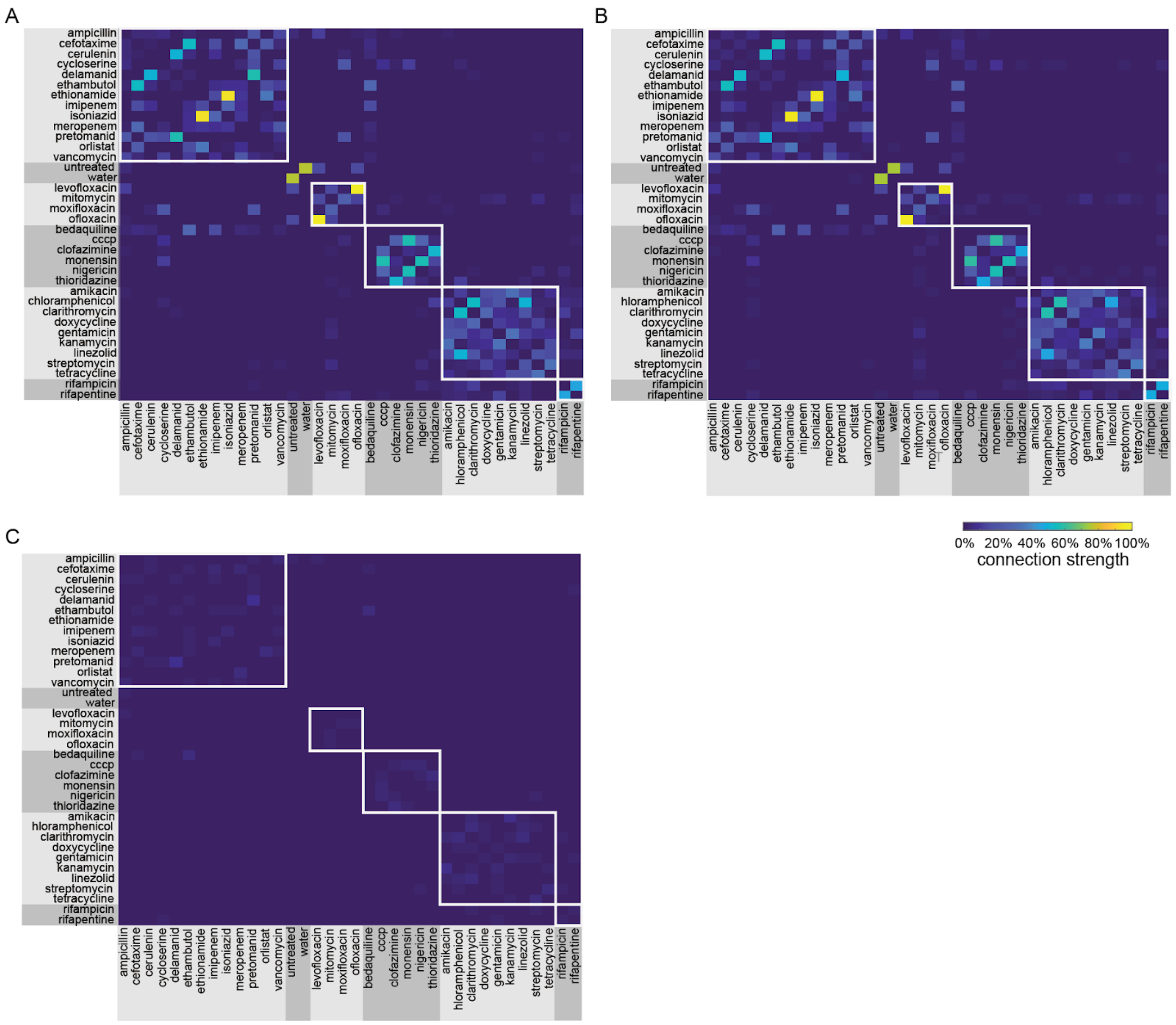


**Fig. S9. 70 classification trials are sufficient for cKNN convergence.** cKNN matrices with 70 (A, also shown in Fig. 3B) and 140 (B) classification trials. The magnitude of the difference in connection strengths are shown in (C).

**Table S1**. **Compounds used in this study.** Primary drug targets, off-target effects, and broad categorizations based on published studies. We determined IC90 values as the minimal concentration of drug needed to inhibit at least 90% of growth. The broad categorization of each drug was assigned based on literature review.

| Drug name | MIC90 (μg/ml) | Solvent | Vendor | Catalog number | Primary target | Off target effects | Reference |
|---|---|---|---|---|---|---|---|
| **CELL WALL SYNTHESIS** | | | | | | | |
| Meropenem | 25 | DMSO | SigmaAldrich | 1392454 | Peptidoglycan (beta lactam) | ATP burst | (4) |
| Ampicillin | 25 | DMSO | SigmaAldrich | A9393 | Peptidoglycan (beta lactam) | | (5) |
| Cefotaxime | 25 | Water | SigmaAldrich | C7039 | Peptidoglycan (beta lactam) | | (5) |
| Isoniazid | 0.047 | DMSO | SigmaAldrich | I3377 | FASII/mycolic acid synthesis (InhA) | ATP burst | (5, 6) |
| Ethambutol | 1.5 | DMSO | Alfa Aesar | J60695 | Arabinogalactan (AftB,AftC,AftD,EmbC) | ATP burst | (5, 6) |

| | | | | | | | |
|---|---|---|---|---|---|---|---|
| Ethionamide | 3.125 | DMSO | TCI Chemicals | E0695 | FASII/mycolic acid synthesis (InhA) | | (5) |
| Imipenem | 3.125 | Water | SigmaAldrich | PHR1796 | Peptidoglycan (beta lactam) | | (5) |
| Vancomycin | 25 | Water | SigmaAldrich | V0045000 | Peptidoglycan (d-ala-d crosslinking) | | (5) |
| Cycloserine | 6.25 | DMSO | Calbiochem | 239831 | Peptidoglycan (d-ala-d ligase) | | (7) |
| Delamanid | 1.5 | DMSO | Advanced ChemBlocks Inc | L13485 | Respiratory toxicity/mycobacterial cell wall (Nitroimidazole) | | (8-10) |
| Pretomanid | 12.5 | DMSO | ApexBio Technology | A1736 | Respiratory toxicity/mycobacterial cell wall (Nitroimidazole) | NO release | (8, 10, 11) |
| Cerulenin | 12.5 | DMSO | SigmaAldrich | C2389 | FASII (KasA, KasB) | | (5) |
| Orlistat | 50 □M* | DMSO | VWR | 89149-186 | PDIM | | (12) |

| | | | | | | | |
|---|---|---|---|---|---|---|---|
| **DNA SYNTHESIS** | | | | | | | |
| Levofloxacin | 3.125 | DMSO | SigmaAldrich | 28266 | DNA gyrase | | (13) |
| Moxifloxacin | 6.25 | DMSO | Alfa Aesa | J66626 | DNA gyrase | | (14) |
| Mitomycin | 12.5 | DMSO | SigmaAldrich | Y0000378 | Alkylation of DNA | Redox recycling | (15) |
| Ofloxacin | 12.5 | 1N NaOH | SigmaAldrich | O8757 | DNA gyrase | | (13) |
| **PROTEIN SYNTHESIS** | | | | | | | |
| Kanamycin | 12.5 | Water | VWR | 408 | 30S subunit of ribosome (irreversible binding); Another source says A-site of 16S rRNA | | (16) |
| Amikacin | 6.25 | Water | SigmaAldrich | A3650 | 30S subunit of ribosome (irreversible binding) | | (16) |
| Chloramphenicol | 25 | DMSO | SigmaAldrich | C0378 | Binds to 23S rRNA of the 50S ribosomal subunit | | (17) |

| | | | | | | | |
|---|---|---|---|---|---|---|---|
| Clarithromycin | 12.5 | DMSO | SigmaAldrich | A3487 | 50S subunit of ribosome | | (18) |
| Doxycycline | 12.5 | DMSO | SigmaAldrich | D9891 | Binds 30S ribosomal subunit | | (19) |
| Gentamycin | 12.5 | Water | SigmaAldrich | G1264 | Aminoglycoside; binds to 30S ribosomal subunit | | (16) |
| Streptomycin | 3.125 | Water | SigmaAldrich | S6501 | Binds to 16S rRNA of the 30S subunit (preventing tRNA from binding to 30S) | | (20, 21) |
| Tetracycline | 50 | DMSO | SigmaAldrich | 87128 | Binds 30S ribosomal subunit (preventing tRNA from binding to 30S) | | (19) |

| | | | | | | | |
|---|---|---|---|---|---|---|---|
| Linezolid | 3.125 | DMSO | ApexBio Technology | A5181 | Binds 50S subunit (prevents 50S from complexing w/ the 30S subunit, mRNA and tRNA) | | (22) |
| **RESPIRATION** | | | | | | | |
| Carbonyl cyanide 3-chlorophenyl-hydrazone | 6.25 | DMSO | SigmaAldrich | C2759 | Proton motive force | | (23) |
| Monensin | 12.5 | MeOH | SigmaAldrich | M5273 | Proton motive force | | (24) |
| Nigericin | 6.25 | MeOH | SigmaAldrich | N7143 | Proton motive force | | (25) |
| Thioridazine | 25 | DMSO | Enzo Life Sciences | BML-NS835-0005 | Electron transport chain - NADH dehydrogenase (NDH-2) | | (25, 26) |
| Pyrazinamide | 50* | DMSO | TCI Chemicals | P0633 | Proton motive force | | (26) |

| | | | | | | | |
|---|---|---|---|---|---|---|---|
| Clofazimine | 6.25 | DMSO | SigmaAldrich | C8895 | Electron transport chain - NDH-2 | | (26, 27) |
| Bedaquiline | 3.125 | DMSO | SigmaAldrich | 10288-25MG | ATP synthase | | (26) |
| **RNAP** | | | | | | | |
| Rifapentine | 0.09 | DMSO | ApexBio Technology | B2127 | RpoB | | (28) |
| Rifampicin | 0.09 | DMSO | TCI Chemicals | R0079 | RpoB | | (26) |
| **Blinded Compounds** | | | | | | | |
| DG167 | 0.39 μM | DMSO | N/A | N/A | KasA | | (29) |
| JSF-3285 | 0.2 μM | DMSO | N/A | N/A | KasA | | Cell Chem. Biol, under revision |
| JSF-2019 | 0.15 μM | DMSO | N/A | N/A | InhA (FAS-II) | | (30) |

*Drugs that did not reach IC90

**Table S2. Features included in analysis pipeline.** For each feature, with exception to FEATURE_1_count and FEATURE_2_count (noted in blue), the median, quartile 1 (25%), quartile 3 (75%), and interquartile range were included as metrics, resulting in 94 total features. Descriptions come from the ImageJ documentation (31).

| Feature | Description | Unit |
|---|---|---|
| FEATURE_1_count | number of times given cell fluoresced with SYTO 24 stain | number of instances |
| FEATURE_2_count | number of FM4-64FX stain positive foci | number of instances |
| **SHAPE_area** | area of cell | $\mu m^2$ |
| **SHAPE_aspectRatio** | major axis/minor axis of cell | unitless |
| **SHAPE_circularity** | 4*[area][perimeter]2 of cell<br><br>ranges from 0 (infinitely elongated polygon) to 1 (perfect circle) | unitless |
| **SHAPE_length** | length of cell | $\mu m$ |
| **SHAPE_perimeter** | perimeter of cell | $\mu m$ |
| **SHAPE_solidity** | [area][convex area] of cell | unitless |
| **SHAPE_width** | width of cell | $\mu m$ |

| | | |
|---|---|---|
| **f_INTENSITY** | intensity of FM4-64FX stain | relative fluorescence units (RFU) |
| **f_SHAPE_area** | area of FM4-64FX foci | $\mu m^2$ |
| **f_SHAPE_aspectRatio** | major axis/minor axis of FM4-64FX foci | unitless |
| **f_SHAPE_circularity** | 4*[area][perimeter]2 of FM4-64FX foci<br><br>ranges from 0 (infinitely elongated polygon) to 1 (perfect circle) | unitless |
| **f_SHAPE_length** | length of FM4-64FX foci | $\mu m$ |
| **f_SHAPE_perimeter** | perimeter of FM4-64FX foci | $\mu m$ |
| **f_SHAPE_solidity** | [area][convex area] of FM4-64FX foci | unitless |
| **f_SHAPE_width** | width of FM4-64FX foci | $\mu m$ |
| **s_INTENSITY** | Intensity of SYTO 24 stain | relative fluorescence units (RFU) |
| **s_SHAPE_area** | area of SYTO 24 foci | $\mu m^2$ |
| **s_SHAPE_aspectRatio** | major axis/minor axis of SYTO 24 foci | unitless |
| **s_SHAPE_circularity** | 4*[area][perimeter]2 of SYTO 24 foci | unitless |

| | | |
|---|---|---|
| | Ranges from 0 (infinitely elongated polygon) to 1 (perfect circle) | |
| **s_SHAPE_length** | length of SYTO 24 foci | μm |
| **s_SHAPE_perimeter** | perimeter of SYTO 24 foci | μm |
| **s_SHAPE_solidity** | [area][convex area] of SYTO 24 foci | unitless |
| **s_SHAPE_width** | width of SYTO 24 foci | μm |

**Movie S1. Time-lapse imaging of *M. smegmatis* before, during, and after treatment with ethambutol.** RpoB-GFP reporter *M. smegmatis* was imaged in nutrient-rich growth conditions in a constant-flow microfluidic device for 10 hours with no antibiotics, followed by a 6-hour drug treatment with 9.375μg/ml of ethambutol, and followed by a 10-hour no drug recovery period.

**Movie S2. Time-lapse imaging of *M. smegmatis* before, during, and after treatment with rifampicin.** RpoB-GFP reporter *M. smegmatis* was imaged in nutrient-rich growth conditions in a constant-flow microfluidic device for 10 hours with no antibiotics, following by a 6-hour drug treatment with 75μg/ml of rifampicin, and followed by a 10-hour no drug recovery period.

**Movie S3. Time-lapse imaging of *M. smegmatis* before, during, and after treatment with moxifloxacin.** RpoB-GFP reporter *M. smegmatis* was imaged in nutrient-rich growth conditions in a constant-flow microfluidic device for 10 hours with no antibiotics, following by a 6-hour drug treatment with 0.781μg/ml of moxifloxacin, and followed by a 10-hour no drug recovery period.
